# Genetic Variation and Disease Severity of Respiratory Syncytial Viruses

**DOI:** 10.1101/2022.02.28.482432

**Authors:** Christopher S. Anderson, Yun Zhang, Anthony Corbett, Chinyi Chu, Alexander Grier, Lu Wang, Xing Qiu, Mathew Mccall, David J. Topham, Edward E. Walsh, Thomas J. Mariani, Richard Scheuermann, Mary T. Caserta

## Abstract

Respiratory Syncytial Virus (RSV) disease in newborns ranges from mild symptoms to severe disease requiring hospitalization. RSV is classified into two subtypes (RSVA and RSVB) based on antigenic and genetic differences. The role these genomic variations play in disease severity remains unknown. Genome sequences were obtained using next-generation RNA sequencing on archived frozen nasal swabs of young children (< 8 months-old) infected with RSV in Rochester, NY between 1977-1998. Samples were chosen from both children hospitalized with severe RSV disease (inpatient) and those presenting with mild symptoms (outpatient) during their first cold-season. Both A and B subtypes demonstrated significant differences in the phylogeny and primary-protein structure during this time period. We found a significant association between RSV phylogeny over this time period and disease severity. For both subtypes, the G-protein demonstrated the greatest amino acid substitutions, although the number of amino acid substitutions was higher in the RSVA subtype. We found a significant association between G-protein variation and disease severity for RSVA, but not RSVB. For both subtypes, variation in the M2-2 protein was significantly associated with disease severity. These results suggest that the genetic variability of RSV proteins may contribute to disease severity in humans.

**Importance:** Each cold-season Respiratory Syncytial Virus (RSV) infects thousands of children in the US. Some will display mild cold symptoms while others develop severe disease, sometimes resulting in lifelong lung problems or fatality. RSV initiates infection and replicates in the nasopharynx. Substitutions in the RSV genome can be found in clinically isolated nasal-swab samples of RSV infected children. Whether these genome variations contribute to severe disease is unknown. Here we found a statistically significant association between RSV phylogeny and disease severity. Furthermore, we found specific RSV proteins (G and M2-2) whose amino acid variation was statistically associated with severe disease, although which protein was associated depended on subtype. Taken together, our results suggest that RSV genotype contributed to disease severity over this time period.

## Introduction

RSV is the leading cause of severe disease in young children. Viral infection occurs primarily in ciliated epithelial cells lining the human airways[1,2]. Acquisition of the virus occurs through either inhalation of aerosolized virus particles or direct contact of the airway epithelial cells with the virus, usually from on our hands[3]. Infection usually presents clinically as a mild respiratory disease with symptoms of rhinitis and cough being most common. For some individuals, especially young children during their first cold season, the virus presents as severe disease with severe fever, cough, and wheeze leading to significant morbidity and sometimes death[4,5]. Long-term effects of severe early-life RSV disease have also been reported[6,7].

The genomes of circulating RSV viruses are not the same and can be grouped into two subtypes (RSVA and RSVB) [3]. Within both subtypes significant genetic variation has occurred over time[8–12]. Moreover, a number of positively selected sites have been identified suggesting the variation is not random but an adaption to external pressure[8].

Most studies have focused on comparisons between RSV subtypes in relation to disease severity, with multiple studies demonstrating increased severity with RSVA, although these studies have been inconsistent[13]. Specific mutations such as those found in the F protein have been shown to result in differences in RSV severity in mice[14]. Furthermore, recent studies of the newly emerged RSV strains have demonstrated differences both in vitro and in vivo[15].

Many methods have been developed in order to statistically associate species variability and phenotype. Whole-viral-genome phylogenetics and phenotype (e.g. disease severity of host) can be statistically compared using a Bayesian association of phylogenetic topologies and phenotypes groups[16]. Additionally, non-parametric, distance-based methods that associate a trait with species diversity, including genetic diversity, have been developed using permutation methods[17,18]. These methods test the homogeneity of dispersion among groups or whether composition among groups are similar[19–21]. Together, these methods provide means to associate RSV genetic variability with disease severity.

## Methods

### Sample Collection

Medical record data was used to identify nasal swab samples positive for RSV by PCR from children hospitalized for severe RSV disease and children seen in outpatient clinics presenting as mild disease. Clinical data, including age at time of infection, was also collected. Original sample collection occurred in the Rochester, NY area from 1977-1998. Nasal swabs were frozen and stored at −80C. Frozen samples were thawed and immediately lyzed with RNA lysis buffer. An RNA sequencing library was prepared and sequenced and 160 samples with whole-genome were obtained.

### Phylogenetics Analysis and Trait Association

Full RSV genomes were aligned using MAFFT. Phylogenetic analysis and trees were produced using RaXML 1000. Bayesian Tip-association Significance testing (BaTS) was performed using the XXX software. The BaTS methods depend on tree topology and use bootstrap replicate trees. The BaTS algorithm applies three statistical methods to test the association between phylogeny and a trait (parsimony score, association index, and maximum exclusive single-state clade size).

### Primary Protein Structure Analysis and Trait Association

Protein peptide sequences were translated in silico from each of the 11 protein-coding-regions for each RSV genome. Protein sequences were aligned using MUSCLE[22]. Pairwise Hamming distances between all aligned sequences were determine using the “stringdist” package in R version 3.4.4. Statistical relationships between the primary protein structures Hamming distance matrix and disease severity phenotype were determined using two statistical approaches. The first statistical method used to determine statistically significant differences between “outpatient” and “inpatient” derived RSV strains was a multivariate test of location in the Hamming distance matrix using the *adonis2* function from the *Vegan* package in R version 3.4.4. 9999 permutations were performed to determine empirical null.

The second approach used a similar method, *anosim* function from the *Vegan* package in R version 3.4.4., but has been reported to be less effected by limited degrees of freedom. 9999 permutations were performed to determine empirical null.

### Association with Disease Severity and Amino Acid Substitutions at each Residue

We used the meta-CATS pipeline [23] to identify statistically significant amino acid positions of RSVA or RSVB subtypes with disease severity status (mild/severe). At each position a chi-square test of independence and Pearson’s chi-square test is performed to calculate a *p*-value.

## Results

RSV viruses were sequenced from nasal swab samples obtained from young children infected in Rochester, New York between 1977 and 1998. Samples were chosen to represent roughly equal sex (42% Female and 58% male). Samples were obtained from children in order to equally represent both mild (outpatient; 87/160 (54%)) and severe (inpatient; 73/160 46%) disease. Samples were chosen to enrich for primary infection sequences by choosing samples of subject that were infected between 0 and 0.8 months old (single “cold season”). PCR-based RSV subtyping data was used in attempt to equally represent A and B subtypes (RSVA = 58% and RSVB = 42%). The number of samples varied year to year (2 - 14 samples per year) with an average of 7.27 samples per year over the 21-year time frame.

Phylogenetic analysis of RSV strains from 1977 until 1998 separated into two distinct linages corresponding to the RSV A and B subtypes (Figure 1). Using a Bayesian approach to phylogenetic association (BEAST), we found a very high consensus on topology. The BaTS algorithm was used to determine if any association between phylogenetics was associated with disease severity status. Phylogeny-trait association demonstrated significant differences between trait (mild/severe disease) distribution and tree topology (Table 1). Both the association index and parsimony association showed statistically association with trait and phylogeny. Interestingly, the maximum exclusive single-state clade size, which is expected to be larger when tips all share the same trait, were significant for the severe trait, but not mild.

**Figure 1.**
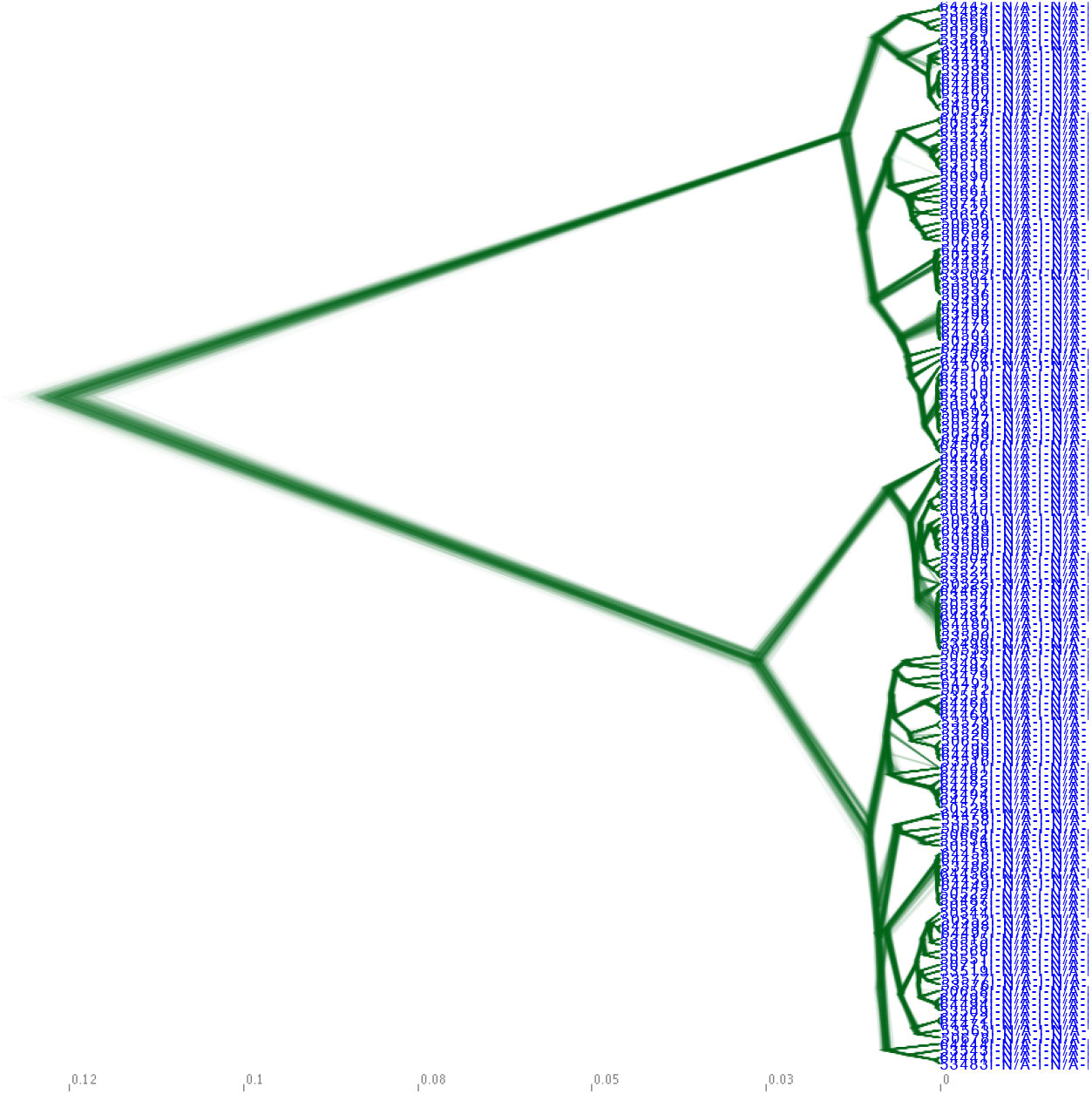
Genomic Variation of the RSV Genome. Aligned whole-genome RSV sequences of viruses collected from nasal swabs of children >8 months-old infected in their first “cold season” between 1977-1998 in Rochester, NY. Phylogenetic trees were fitted using a Bayesian approach (BEAST). To visualize both uncertainty in node heights and uncertainty in topology phylogenetic trees were visualized using DensiTree.

**Table 1.**

| Statistic | observed<br>mean | lower<br>95% CI | upper<br>95% CU | null<br>mean | lower<br>95% CI | upper<br>95% CI | significanc<br>e |
| --- | --- | --- | --- | --- | --- | --- | --- |
| AI | 7.025 | 6.305 | 7.752 | 8.862 | 7.387 | 10.371 | 0.023 |
| PS | 45.935 | 44.000 | 47.000 | 53.798 | 48.308 | 58.778 | 0.012 |
| MC (Severe) | 9.000 | 9.000 | 9.000 | 4.809 | 3.153 | 7.233 | 0.027 |
| MC (Mild) | 4.022 | 4.000 | 4.000 | 3.896 | 2.632 | 6.000 | 0.503 |

Of the 11 RSV proteins, the G protein, for both subtypes, showed the maximum number of total amino acid substitutions (RSVA G protein = 64, RSVB G protein = 53; Figure 2A) as well as the greatest percent change per amino acid length of any protein (RSVA G protein = 21%, RSVB G protein = 17%; Figure 2B). The M2-2 protein was also one of the most variable proteins both in the number of total amino acids (RSVA M2-2 protein = 16, RSVB M2-2 protein = 11) and percent change per protein length (RSVA M2-2 protein = 18%, RSVB M2-2 protein = 12%). The L protein showed many substitutions for both subtypes (RSVA L protein = 43, RSVB L protein = 30) although the per amino acid change was moderate (RSVA L protein = 2%, RSVB L protein = 1%). Alternatively, the SH protein and F protein showed lower numbers of amino acid substitutions (RSVA SH protein = 5, RSVB SH protein = 6; RSVA F protein = 14, RSVB F protein = 10), but SH showed a moderate change per amino acid compared to other proteins (RSVA SH protein = 8%, RSVB SH protein = 9%) and the F protein showed a minimal number of substitutions per protein length (RSVA F protein = 2%, RSVB F protein = 2%). All other proteins showed both minimal substitutions (3 - 6) and percent changes (1 – 4%).

**Figure 2.**
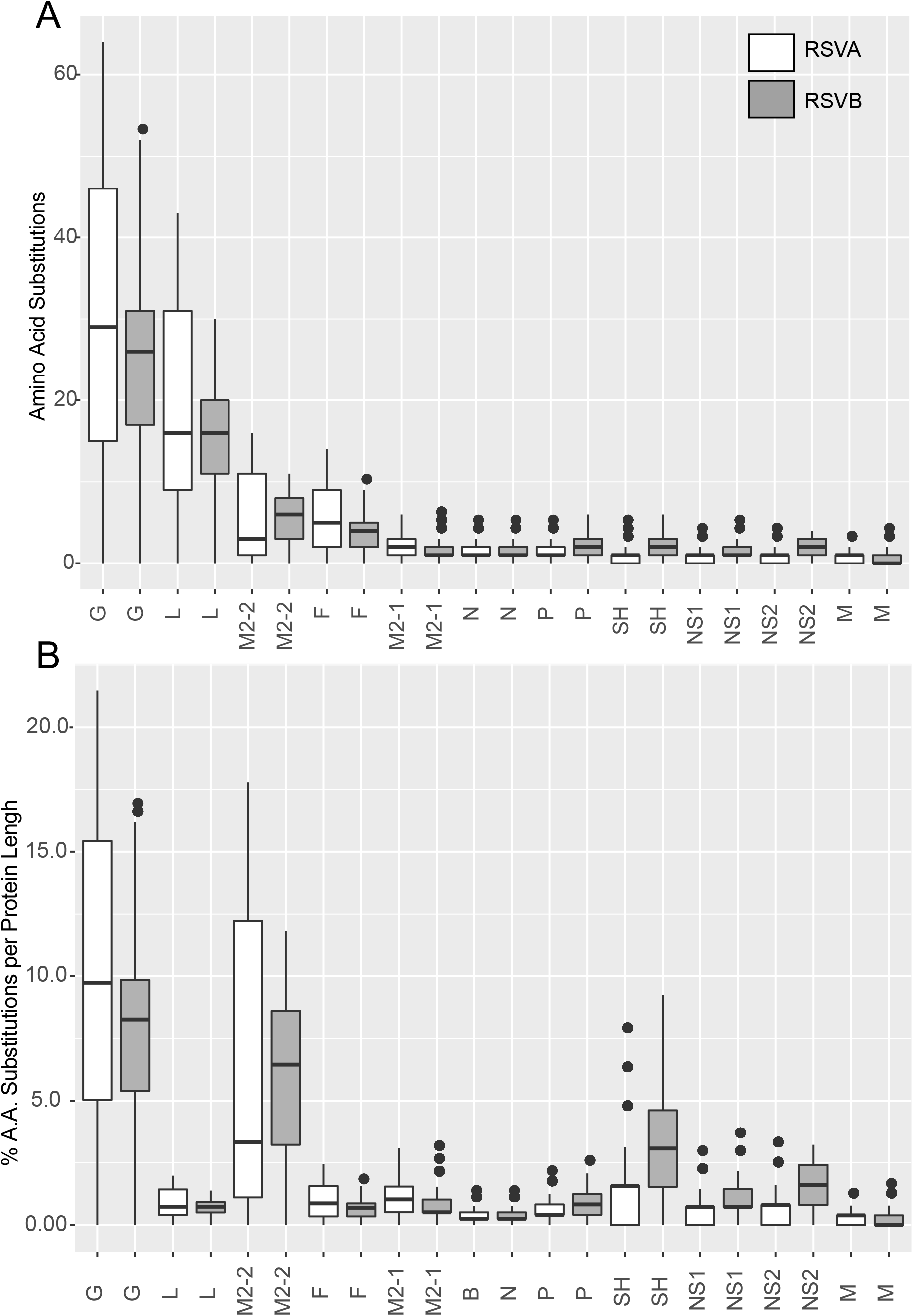
Comparison of Amino Acid Variation in Across RSV Proteins. RSV protein sequences within subtypes were aligned using MUSCLE. The number of amino acid substitution between each RSV sequence was calculated. (A) Boxplot of the number of amino acid substitutions between all RSV proteins by subtype. (B) Boxplot of the percentage of number of amino acid substitutions divided by the amino acid length of the protein between all RSV proteins by subtype.

We next determined if amino acid variation in specific viral proteins were associated with disease severity. Using two permutation-based statistical approaches, we determined if amino acid variability was associated with disease phenotype (mild/severe). We found that in both subtypes the M2-2 protein was significantly associated with disease severity (Table 2; Figure 3). For the RSV A subtype, the G protein was also significantly associated with disease severity. The NS2 protein was also significantly associated with disease severity in the RSVB subtype, although only for one statistical test.

**Figure 3.**
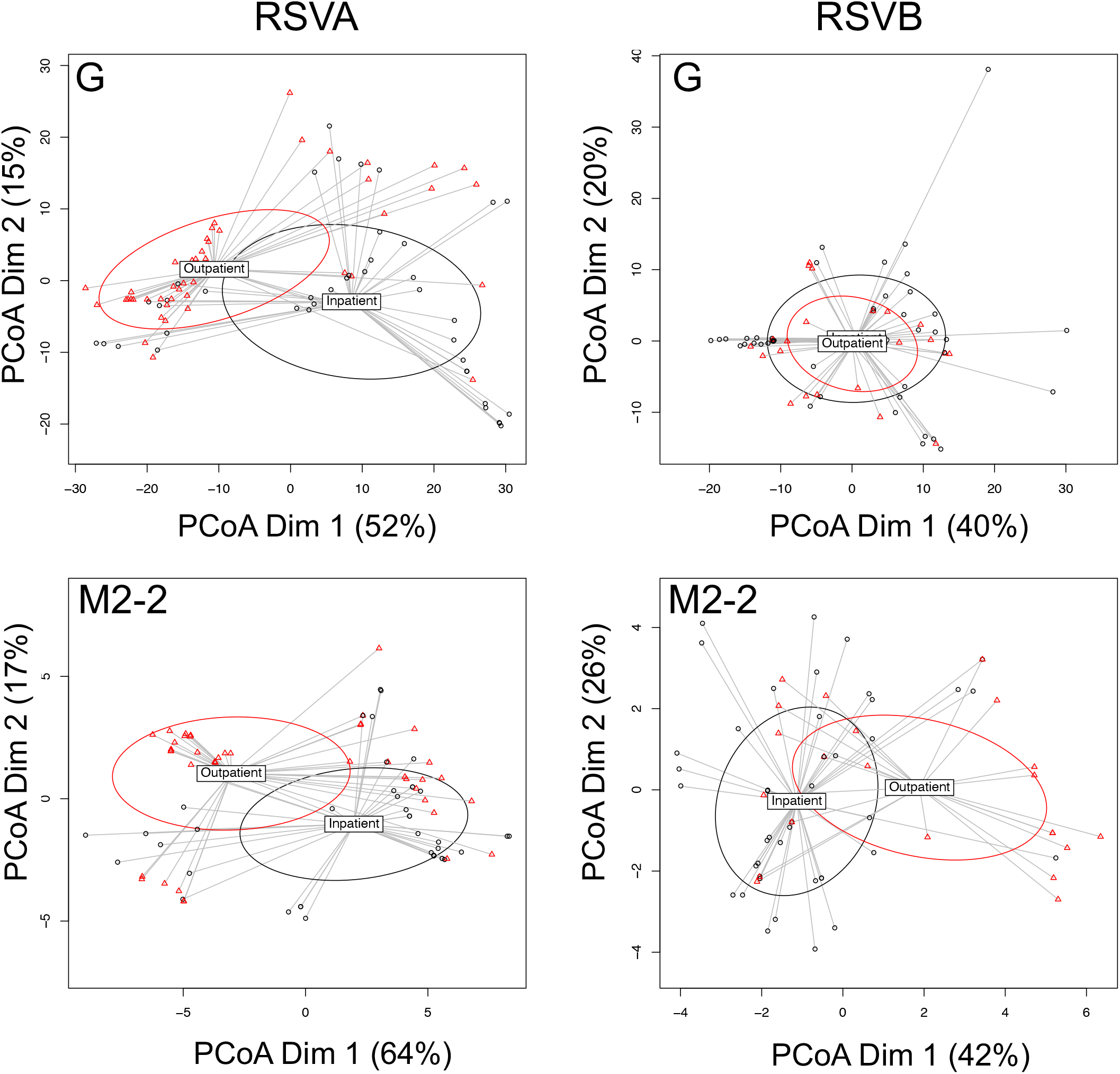
Primary Protein Structure Variability Among RSV G and M2-2 Proteins. Protein sequences for G and M2-2 proteins from RSVA and RSVB subtypes were aligned separately. The number of amino acid substitutions were calculated between all strains resulting. Principal coordinate analysis was performed to demonstrate amino acid variability in reduced dimensional space. Ellipses are centered on centroids with 1 standard deviation. Points are colored by disease severity status; red = mild/outpatient, black = severe/inpatient. When points contain multiple sequences and from patients of both disease types, points are colored by the more numerous disease type.

**Table 2.**

| Protein | Subtype | Anosim<br>Rval | Anosim<br>pval | Anosim<br>pval.adj | Adonis2<br>Fval | Adonis2<br>pval | Adonis2<br>pval.adj |
| --- | --- | --- | --- | --- | --- | --- | --- |
| G | A | 0.122 | 0.001 | 0.011* | 9.438 | 0.001 | 0.011* |
| G | B | -0.140 | 0.999 | 0.999 | -4.832 | 1.000 | 1.000 |
| L | A | 0.026 | 0.068 | 0.166 | 2.916 | 0.071 | 0.137 |
| L | B | -0.005 | 0.515 | 0.629 | 2.083 | 0.081 | 0.137 |
| M2-2 | A | 0.119 | 0.001 | 0.011* | 7.251 | 0.004 | 0.022* |
| M2-2 | B | 0.105 | 0.007 | 0.051* | 5.078 | 0.002 | 0.015* |
| F | A | 0.038 | 0.040 | 0.126 | 2.602 | 0.093 | 0.138 |
| F | B | 0.085 | 0.068 | 0.166 | 3.573 | 0.023 | 0.072 |
| M2-1 | A | 0.014 | 0.153 | 0.259 | 2.843 | 0.073 | 0.137 |
| M2-1 | B | 0.040 | 0.209 | 0.307 | 3.178 | 0.036 | 0.088 |
| N | A | 0.004 | 0.306 | 0.396 | 2.398 | 0.100 | 0.138 |
| N | B | 0.101 | 0.024 | 0.114 | 2.179 | 0.095 | 0.138 |
| P | A | 0.011 | 0.197 | 0.307 | 1.129 | 0.337 | 0.412 |
| P | B | 0.109 | 0.026 | 0.114 | 6.393 | 0.017 | 0.072 |
| SH | A | 0.020 | 0.111 | 0.218 | 2.676 | 0.075 | 0.137 |
| SH | B | -0.079 | 0.975 | 0.999 | 0.327 | 0.765 | 0.842 |
| NS1 | A | 0.041 | 0.036 | 0.126 | 4.224 | 0.026 | 0.072 |
| NS1 | B | -0.066 | 0.940 | 0.999 | 0.097 | 0.885 | 0.927 |
| NS2 | A | 0.020 | 0.119 | 0.218 | 1.148 | 0.321 | 0.412 |
| NS2 | B | -0.027 | 0.695 | 0.805 | 7.075 | 0.001 | 0.011* |
| M | A | 0.021 | 0.091 | 0.200 | 5.191 | 0.021 | 0.072 |
| M | B | 0.022 | 0.303 | 0.396 | 0.244 | 0.721 | 0.835 |

\* p value ≤ 0.05

We sought to evaluate if any specific mutations were significantly associated with disease severity. We compared each residue in the RSVA G and M2-2 proteins and RSVB M2-2 proteins with disease severity. Two out of the three proteins, RSVA G and RSVB M2-2, had significant mutations associated with disease severity (Table 3). RSV A G-protein had seven amino acids associated severity status, while the RSVB M2-2 protein had three amino acids associated. Taken together, our results suggest that certain genetic variations in RSV may be more likely to be seen in viruses isolated from young children hospitalized with RSV.

**Table 3.**

| Subtype | Protein | Position | Chi-square Value | P-value | Degree Freedom | Severe Residue Diversity | Mild Residue Diversity |
| --- | --- | --- | --- | --- | --- | --- | --- |
| A | G | 1528 | 8.913 | 0.0116 | 2 | 7 L, 15 P, 20 S | 5 L, 34 P, 12 S |
| A | G | 1327 | 7.558 | 0.02285 | 2 | 14 L, 4 P, 24 Q | 11 L, 40 Q |
| A | G | 1298 | 4.159 | 0.04142 | 1 | 22 N, 20 S | 15 N, 36 S |
| A | G | 1496 | 8.215 | 0.04176 | 3 | 22 H, 18 L, 2 N | 37 H, 12 L, 2 Y |
| A | G | 1524 | 4.007 | 0.0453 | 1 | 22 L, 20 P | 38 L, 13 P |
| A | G | 1324 | 6.123 | 0.04682 | 2 | 18 I, 4 N, 20 T | 12 I, 2 N, 37 T |
| A | G | 1414 | 3.923 | 0.04764 | 1 | 24 K, 18 R | 40 K, 11 R |
| B | M2-2 | 2341 | 5.57 | 0.01827 | 1 | 30 I, 14 V | 8 I, 15 V |
| B | M2-2 | 2354 | 4.905 | 0.02678 | 1 | 41 H, 3 Y | 16 H, 7 Y |
| B | M2-2 | 2551 | 4.905 | 0.02678 | 1 | 3 D, 41 E | 7 D, 16 E |

## Discussion

We sought to provide insight into RSV-associated severe respiratory disease in young children experiencing their primary infection. Although many host factors are recognized as contributors to severe disease[5], the contribution of the virus genetics has not been well explored. In the study, we assessed genomic variation of RSV viruses that circulated in Rochester, New York from 1977 - 1998. Our findings confirm that the RSV genotype changes over time and multiple genotypes circle each year. Furthermore, our results demonstrate that RSV genetic variation is not spatially restricted and local regions are exposed to a multitude of unique RSV strains. We compared RSV sequence variation and disease severity using both phylogenetic and non-phylogenetic approaches. Phylogenetic approaches demonstrated that both tree topography, including monophyletic clades, were associated with severe disease. Lastly, our results suggest that RSV strains with specific amino acid substitutions in the G or M2-2 proteins contribute to disease severity in young children.

What specific impact amino acid substitutions in the surface proteins of RSV have on the host defense and if these changes result in antigenic drift is still largely unexplored, although the recently emerged ON1 RSVA strain containing a duplication in G has been shown to cause increase severe disease[24]. We found changes in the G protein were the most predominant. Additionally, the SH protein showed minor variation and was not associated with disease severity. The F protein also varied, but was not associated with disease severity during primary infection, although others have demonstrate changes in F that do increase disease severity[14]. Future studies will be needed to better understand the relationship between surface protein mutation and RSV disease.

We were surprised to see the structural protein M2-2 was associated with severity. M2-2 has been shown to be involved in viral RNA transcription and replication regulations. Furthermore, a current vaccine candidate has a M2-2 gene deletion that attenuates the virus, potentially providing protection, but resulting in mild disease. Whether variation in the M2-2 gene effects the transcription/replication regulation process is unknown.

Taken together, our results suggest that RSV variation can impact disease severity. Although our studies were not designed to investigate mechanism or causality, they do suggest that changes in RSV genes are associated with disease severity in the very young experiencing a primary infection. Whether these changes are due to the adaptive immune response, or random genetic drift, is still unknown and future studies will be needed to confirm if variation the RSV genome affects disease severity.

## Acknowledgements

We would like to thank all sample donors for their contribution to the study. The original work was supported by the University of Rochester Pulmonary Training Grant T32-HL066988, and the Health Sciences Center for Computational Innovation Pilot Award OP211341.

## References

[1] C.S. Anderson, C.-Y. Chu, Q. Wang, J.A. Mereness, Y. Ren, K. Donlon, et al., CX3CR1 as a respiratory syncytial virus receptor in pediatric human lung, Pediatr. Res. 87 (2020) 862–867. doi:10.1038/s41390-019-0677-0.

[2] K.-I. Jeong, P.A. Piepenhagen, M. Kishko, J.M. DiNapoli, R.P. Groppo, L. Zhang, et al., CX3CR1 Is Expressed in Differentiated Human Ciliated Airway Cells and Co-Localizes with Respiratory Syncytial Virus on Cilia in a G Protein-Dependent Manner, PLoS ONE. 10 (2015) e0130517. doi:10.1371/journal.pone.0130517.

[3] B.N. Fields, Fields Virology, Stanford University Press, 2013. doi:10.11126/stanford/9780804770750.001.0001.

[4] T. Shi, D.A. McAllister, K.L. O’Brien, E.A.F. Simoes, S.A. Madhi, B.D. Gessner, et al., Global, regional, and national disease burden estimates of acute lower respiratory infections due to respiratory syncytial virus in young children in 2015: a systematic review and modelling study, Lancet. 390 (2017) 946–958. doi:10.1016/S0140-6736(17)30938-8.

[5] M.T. Caserta, X. Qiu, B. Tesini, L. Wang, A. Murphy, A. Corbett, et al., Development of a Global Respiratory Severity Score for Respiratory Syncytial Virus Infection in Infants, J. Infect. Dis. 215 (2017) 750–756. doi:10.1093/infdis/jiw624.

[6] N. Sigurs, P.M. Gustafsson, R. Bjarnason, F. Lundberg, S. Schmidt, F. Sigurbergsson, et al., Severe respiratory syncytial virus bronchiolitis in infancy and asthma and allergy at age 13, Am. J. Respir. Crit. Care Med. 171 (2005) 137–141. doi:10.1164/rccm.200406-730OC.

[7] P. Wu, T.V. Hartert, Evidence for a causal relationship between respiratory syncytial virus infection and asthma, Expert Rev Anti Infect Ther. 9 (2011) 731–745. doi:10.1586/eri.11.92.

[8] A. Trento, M. Viegas, M. Galiano, C. Videla, G. Carballal, A.S. Mistchenko, et al., Natural history of human respiratory syncytial virus inferred from phylogenetic analysis of the attachment (G) glycoprotein with a 60-nucleotide duplication, J. Virol. 80 (2006) 975–984. doi:10.1128/JVI.80.2.975-984.2006.

[9] H. Chi, K.-L. Hsiao, L.-C. Weng, C.-P. Liu, H.-F. Liu, Persistence and continuous evolution of the human respiratory syncytial virus in northern Taiwan for two decades, Sci. Rep. 9 (2019) 4704–9. doi:10.1038/s41598-019-41332-9.

[10] J.A. Melero, M.L. Moore, Influence of respiratory syncytial virus strain differences on pathogenesis and immunity, Curr. Top. Microbiol. Immunol. 372 (2013) 59–82. doi:10.1007/978-3-642-38919-1_3.

[11] W.M. Sullender, Respiratory syncytial virus genetic and antigenic diversity, Clin. Microbiol. Rev. 13 (2000) 1–15– table of contents. doi:10.1128/cmr.13.1.1-15.2000.

[12] L. Tan, F.E.J. Coenjaerts, L. Houspie, M.C. Viveen, G.M. van Bleek, E.J.H.J. Wiertz, et al., The comparative genomics of human respiratory syncytial virus subgroups A and B: genetic variability and molecular evolutionary dynamics, J. Virol. 87 (2013) 8213–8226. doi:10.1128/JVI.03278-12.

[13] S. Vandini, C. Biagi, M. Lanari, Respiratory Syncytial Virus: The Influence of Serotype and Genotype Variability on Clinical Course of Infection, Int J Mol Sci. 18 (2017). doi:10.3390/ijms18081717.

[14] M.L. Moore, M.H. Chi, C. Luongo, N.W. Lukacs, V.V. Polosukhin, M.M. Huckabee, et al., A chimeric A2 strain of respiratory syncytial virus (RSV) with the fusion protein of RSV strain line 19 exhibits enhanced viral load, mucus, and airway dysfunction, J. Virol. 83 (2009) 4185–4194. doi:10.1128/JVI.01853-08.

[15] F. Midulla, G. Di Mattia, R. Nenna, C. Scagnolari, A. Viscido, G. Oliveto, et al., Novel Variants of Respiratory Syncytial Virus A ON1 Associated With Increased Clinical Severity of Bronchiolitis, J. Infect. Dis. 222 (2020) 102–110. doi:10.1093/infdis/jiaa059.

[16] J. Parker, A. Rambaut, O.G. Pybus, Correlating viral phenotypes with phylogeny: accounting for phylogenetic uncertainty, Infect. Genet. Evol. 8 (2008) 239–246. doi:10.1016/j.meegid.2007.08.001.

[17] M.J. Anderson, A new method for non-parametric multivariate analysis of variance, Austral Ecology. 26 (2001) 32–46. doi:10.1111/j.1442-9993.2001.01070.pp.x.

[18] J.E. Bennett, R. Dolin, M.J. Blaser, Mandell, Douglas, and Bennetts. Principles and Practice of Infectious Diseases, Philadelphia: Churchill Livingstone, 2015.

[19] B.H. McArdle, M.J. Anderson, FITTING MULTIVARIATE MODELS TO COMMUNITY DATA: A COMMENT ON DISTANCE-BASED REDUNDANCY ANALYSIS, Ecology. 82 (2001) 290–297. doi:10.1890/0012-9658(2001)082[0290:FMMTCD]2.0.CO;2@10.1002/(ISSN)1939-9170(CAT)VirtualIssues(VI)scECY.

[20] M.J. Anderson, Distance-based tests for homogeneity of multivariate dispersions, Biometrics. 62 (2006) 245–253. doi:10.1111/j.1541-0420.2005.00440.x.

[21] M.J. Anderson, K.E. Ellingsen, B.H. McArdle, Multivariate dispersion as a measure of beta diversity, Ecol. Lett. 9 (2006) 683–693. doi:10.1111/j.1461-0248.2006.00926.x.

[22] R.C. Edgar, MUSCLE: multiple sequence alignment with high accuracy and high throughput, Nucleic Acids Res. 32 (2004) 1792–1797. doi:10.1093/nar/gkh340.

[23] B.E. Pickett, M. Liu, E.L. Sadat, R.B. Squires, J.M. Noronha, S. He, et al., Metadata-driven comparative analysis tool for sequences (meta-CATS): an automated process for identifying significant sequence variations that correlate with virus attributes, Virology. 447 (2013) 45–51. doi:10.1016/j.virol.2013.08.021.

[24] A. Streng, D. Goettler, M. Haerlein, L. Lehmann, K. Ulrich, C. Prifert, et al., Spread and clinical severity of respiratory syncytial virus A genotype ON1 in Germany, 2011-2017, BMC Infect. Dis. 19 (2019) 613–10. doi:10.1186/s12879-019-4266-y.

